# Cultivation-based quantification and identification of bacteria at two hygienic key sides of domestic washing machines

**DOI:** 10.1101/2021.02.19.431940

**Authors:** Susanne Jacksch, Huzefa Zohra, Mirko Weide, Sylvia Schnell, Markus Egert

**Affiliations:** Faculty of Medical and Life Sciences, Institute of Precision Medicine, Microbiology and Hygiene Group, Furtwangen University, 78054 Villingen-Schwenningen, Germany; International Research & Development – Laundry & Home Care, Henkel AG & Co. KGaA, 40191 Düsseldorf, Germany; Institute of Applied Microbiology, Research Centre for BioSystems, Land Use, and Nutrition (IFZ), Justus-Liebig-University Giessen, 35392 Giessen, Germany

**Keywords:** Washing machine, bacteria, hygiene, MALDI Biotyping

## Abstract

Detergent drawer and door seal represent important sites for microbial life in domestic washing machines. Interestingly, quantitative data on microbial contamination of these sites is scarce. Here, 10 domestic washing machines were swap-sampled for subsequent bacterial cultivation at four different sampling sites, each: detergent drawer, detergent drawer chamber as well as top and bottom part of the rubber door seal. The average bacterial load over all washing machines and sites was 2.1 ± 1.0 × 10^4^ CFU cm^−2^ (average ± standard error of the mean (SEM)). The top part of the door seal showed the lowest contamination (11.1 ± 9.2 × 10^1^ CFU cm^−2^), probably due to less humidity.

Out of 212 isolates, 178 (84%) were identified on genus level and 118 (56%) on species level using MALDI biotyping, resulting in 29 genera and 40 identified species across all machines. The predominant bacterial genera were *Staphylococcus* and *Micrococcus*, which were found at all sites. 21 out of 40 species were classified as opportunistic pathogens, emphasizing the need for regular cleaning of the investigated sites.

## Introduction

Representing wet, warm and nutrient-rich environments, many sites of domestic washing machines offer ideal living conditions for microorganisms, such as bacteria and fungi (1, 2). Microbial contamination of washing machines might cause unaesthetic stainings as well as malodor formation (3, 4). In addition, microbial biofilms might serve as reservoirs for (potentially) pathogenic microorganisms that might contaminate the laundry and thereby pose a health threat for susceptible persons (5, 6).

Various studies have shown that washing machines are colonized by a considerable diversity of microbes, often capable of forming biofilms (4, 7–10). For instance, Nix and co- workers (10) investigated pro- and eukaryotic microorganisms on the rubber door seal and the detergent drawer using 16S rRNA gene and ITS1 region pyrosequencing. They identified *Proteobacteria* as the main bacterial representatives and *Basidiomycota* and *Ascomycota* as the main fungal colonizers (10).

Regarding bacteria, washing machines are indeed mainly populated by the phyla *Proteobacteria, Actinobacteria, Firmicutes* and *Bacteroidetes* (7, 9, 10). They largely enter the machine via soiled clothing, tap water, and maybe also air (2, 4). In a recent molecular study on the bacterial community of domestic washing machines, we identified the detergent drawer as the site with the highest bacterial diversity and the door seal as the site with highest relative abundance of malodor forming *Moraxella osloensis* species. Per site, 30-60% of the relatively most abundant sequence types were closely related to potentially pathogenic bacteria, such as *Brevundimonas vesicularis* or *Pseudomonas aeruginosa* inside the detergent drawer and *Moraxella osloensis* or *Acinetobacter parvus* inside the door seal (9). In a startling study, an antibiotic resistant *Klebsiella oxytoca* strain was recently isolated from biofilms of detergent drawer and door seal of a domestic washing machine used for the woollen laundry of a paediatric hospital ward, from which it probably had colonized newborns (11).

Interestingly, quantitative data on the microbial contamination of different sites of domestic washing machines is scarce. To increase knowledge in this field, we aerobically cultivated and quantified bacteria from two sites of the detergent drawer and the door seal region of 10 domestic washing machines, each, and identified representative isolates by MALDI Biotyping.

## Material and Methods

### Washing machine sampling

Swab samples were taken from 10 domestic, front loading washing machines in the greater area of Villingen-Schwenningen, Germany, between April and June 2020. Each machine was sampled at the detergent drawer, the detergent drawer chamber, and the top and bottom part of the rubber door seal. All tested machines were provided voluntarily by their owners. Similar sites of each machine were swapped with sterile cotton swaps (Deltalab, Rubí, Spain) pre-moistened in sterile physiological (0.9%) saline solution. The sampling area was ~42 cm^2^ for the detergent drawer, ~28 cm^2^ for the detergent chamber and ~45 cm^2^ for the upper and lower parts of the rubber door seal, respectively. After sampling, the swab heads were transferred to a sterile reaction tube containing 2 ml of sterile physiological saline solution. All samples were processed within 1 h after sampling.

### Colony counting

Swab heads were vortexed for 1 min at maximum speed. After serial decimal dilution up to 10^−6^ with sterile physiological saline solution, 100 μl of each dilution were spread in duplicates on tryptic soy agar plates (TSA; Carl Roth Karlsruhe, Germany) and incubated under aerobic conditions for 48 h at 37°C. Subsequently, colonies in the range of 3 - 300 colonies were counted, averaged, and used for calculation of microbial loads per cm^2^ of sample area.

One representative of each colony morphotype (differing in size, color and/or colony morphology) per sample was picked with a sterile inoculation loop, re-streaked on TSA and incubated aerobically at 37°C. After control for purity, a colony from each morphotype was selected, dissolved in 300 μl of MALDI water (Honeywell, Offenbach, Germany) and stored at −80°C for subsequent identification by Matrix-Assisted Laser Desorption/Ionization (MALDI) Biotyping.

### Identification of isolates by MALDI Biotyping

The obtained isolates were identified with a MALDI Biotyper Microflex system (Bruker Daltonics, Bremen, Germany) according to the manufacturer’s instructions. The protein extraction method was applied using ethanol/formic acid sample preparation. 1 μl of the respective protein extract of each isolated colony was added to a spot on the Biotyper steel target plate. After air drying, the samples were overlayed with 1 μl MALDI-matrix solution (alpha-Cyano-4-hydroxycinnamic acid, Bruker Daltonics). After further air drying, the samples were analyzed. The obtained mass spectra were compared against the internal MALDI Biotyper reference libraries: MBT Compass Library, revision F, version 9, containing 8468 main spectra (MSPs); MBT Filamentous Fungi Library (revision No. 2, containing 468 MSPs); MBT Security Related Library (SR Library, revision No. 1; containing 104 MSPs). Matches with the respective spectra in the databases were displayed as scores ranging from 0.0 – 3.0. Scores ≥ 1.7 indicated a secure genus identification, scores ≥ 2.0 a secure genus and probable species identification (12).

### Statistical analyses

The statistical analysis was performed using R (version 3.6.1) (13) and R Studio (version 1.2.1335) (14) with the packages ggplot2 (version 3.2.1) (15), reshape2 (version 1.4.3) (16), and scales (version 1.0.0) (17). Non-parametric tests (Kruskal-Wallis rank sum test followed by Wilcoxon-Mann-Whitney post-hoc tests) were used to check for statistical significance between the colony counts of the four sampling sites. P-values < 0.05 were considered as statistically significant.

## Results and Discussion

### Colony counts at the different sampling sites

All investigated samples showed microbial growth. Microbial loads spanned five orders of magnitude (Figure 1). The average colony count over all samples was 2.1 ± 1.0 × 10^4^ CFU cm^−2^ (average ± standard error of the mean (SEM)). The sampling site with the lowest cell numbers was the top part of the rubber door seal (RDST 11.1 ± 9.2 × 10^1^ CFU cm^−2^), probably because water quickly drains off from here. Detergent drawer (DD), detergent drawer chamber (DC) and bottom part of the rubber door seal (RDSB) showed similar values with 1.1 ± 0.74 × 10^4^ CFU cm^−2^, 4.2 ± 3.0 × 10^4^ CFU cm^−2^ and 3.1 ± 1.9 × 10^4^ CFU cm^−2^, respectively. Statistical analysis proved a significant difference when comparing the colony counts of all sampling sites (p = 0.029, Figure 1). Subsequent pair-wise post-hoc tests indicated differences between the top part of the rubber seal and its bottom part (p = 0.007) as well as the detergent drawer (p = 0.021) and the detergent drawer chamber (p = 0.045). Clearly, due to the large variability of the colony counts, studies with larger sample sizes are needed to substantiate these findings.

**Figure 1.**
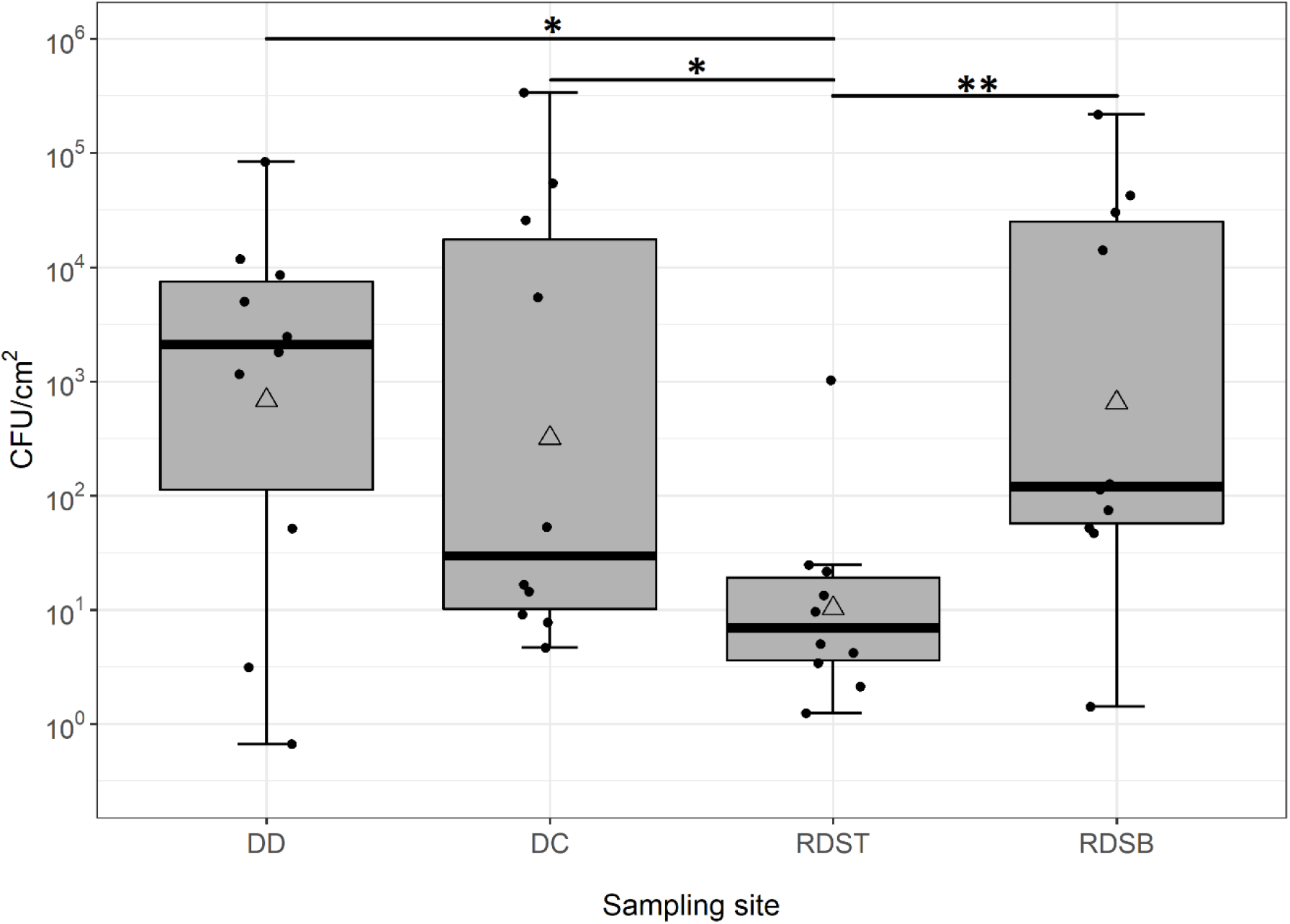
Box-whisker plots of aerobic colony counts per square cm from 4 sampling sites of 10 domestic washing machine. Each box represents the 25% and 75% percentiles. Bold horizontal lines represents medians. Mean values are displayed as triangles. Whiskers above and below the boxes indicate the lowest and highest microbial counts, which were not classified as outliers. Black points represent single data points per site. The different sampling sites are detergent drawer (DD), detergent drawer chamber (DC), top part of rubber door seal (RDST) and bottom part of rubber door seal (RDSB) (n=10 for each sampling site). Results of statistical tests are discussed in the text; significance levels are indicated by asterisks (* = p < 0.05; ** p = < 0.01).

Interestingly, little is known about the microbial load of different sites inside domestic washing machines (18). To the best of our knowledge, only Stapelton and colleagues (18) have previously reported microbial loads of different sampling sites, albeit only for 4 domestic washing machines. While our data match their results for the rubber doors seal quite well (~ 10^3^ to 10^4^ cm^−2^), they also suggest the detergent drawer region to be significantly more contaminated than reported by them (~ 10^−1^ to 10^3^ cm^−2^). Clearly, also from a quantitative point for view, the detergent drawer region is an important site for washing machine hygiene and thus probably also laundry hygiene.

### Identification of microbial isolates

212 microbial isolates stemming from the 40 washing machine samples were analyzed by MALDI Biotyping. Genus-level identifications scores (≥ 1.7) were determined for 178 isolates (84%), while 34 isolates (16%) could not be identified. 118 isolates (56%) were probably identified on species level (score ≥ 2.0). In total, 29 genera and 40 species were found (Table 1).

**Table 1.**
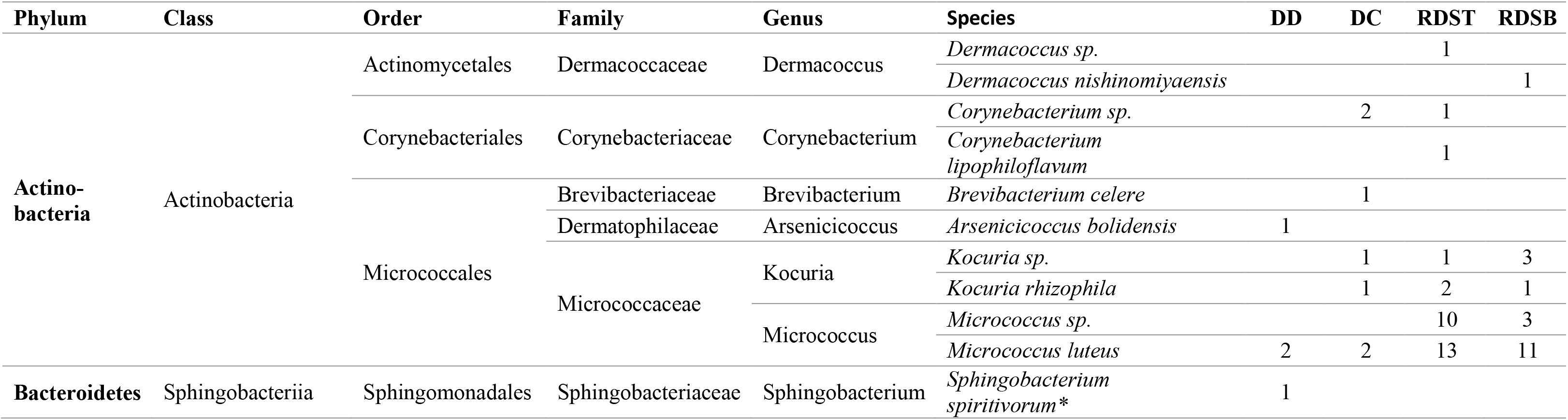
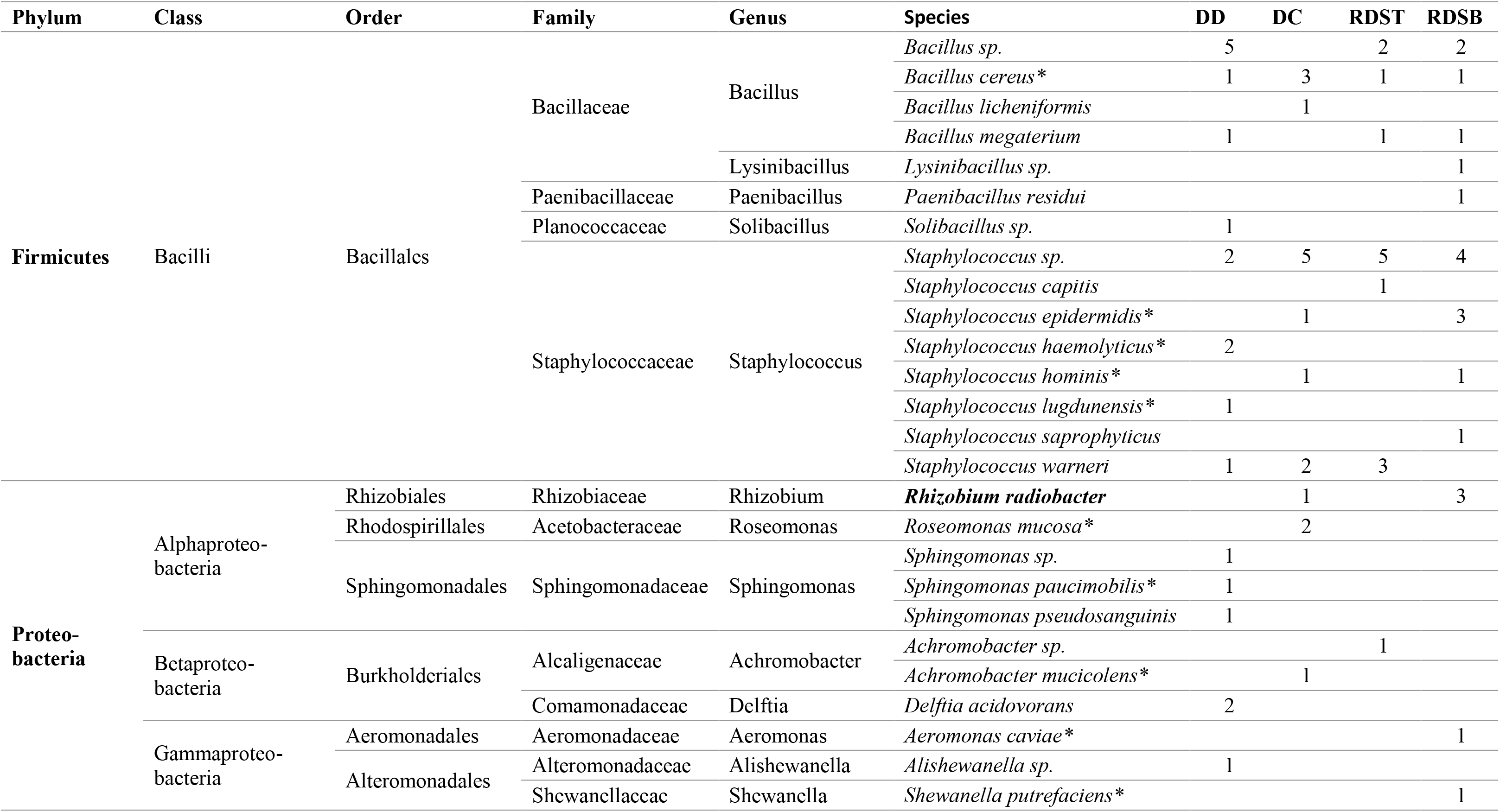
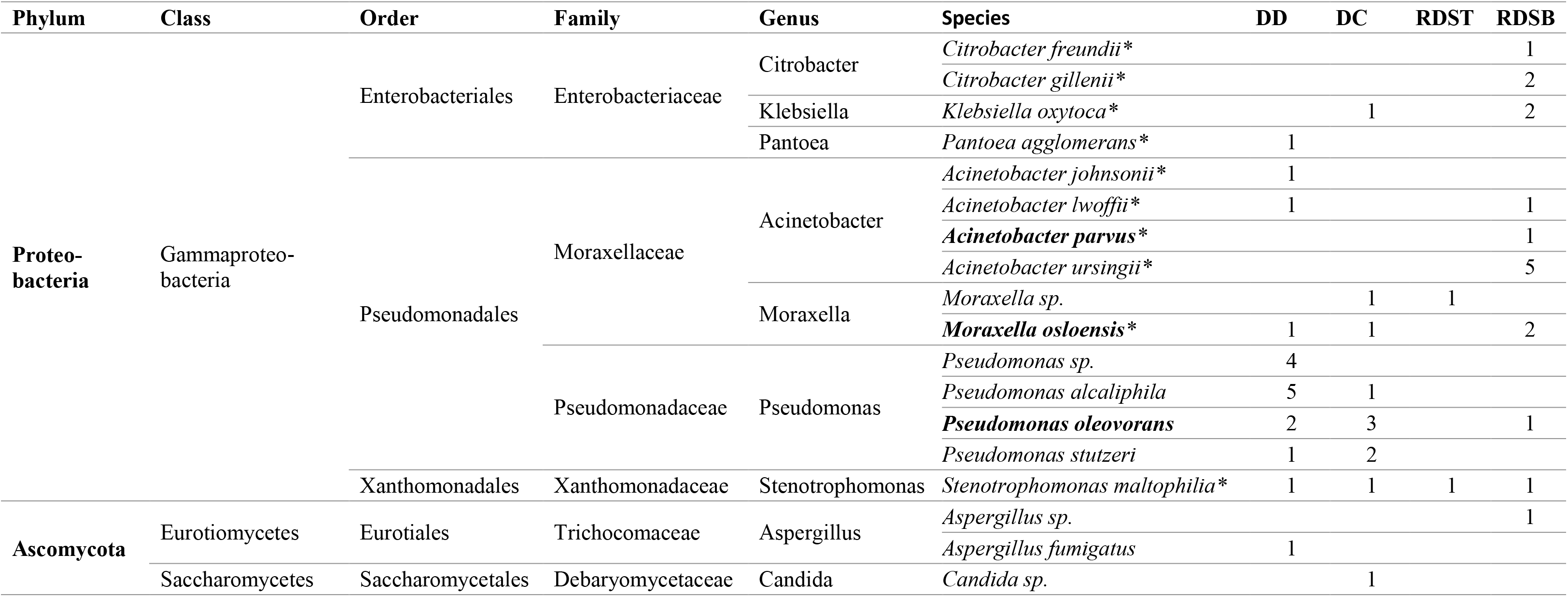
Number of microbial isolates obtained from 10 domestic washing machines and identified by MALDI biotyping with identification scores ≥.1.7 (genus level; n= 178) and ≥ 2.0 (species level; n=118) across the four different sampling sites (DD = detergent drawer; DC = detergent drawer chamber; RDST = top part of rubber door seal; RDSB= bottom part of rubber door seal). Species categorized as risk group 2 (BAuA 2015 (23)) are marked with an asterisk. Species detected here which have been previously identified in (9) as one of the ten relatively most abundant OTUs from door seal and the detergent drawer, respectively, are written in bold.

In accordance with previous studies (7, 9, 10), *Proteobacteria* (29%), *Actinobacteria* (27%), *Firmicutes* (26%) and *Bacteroidetes* (0.5%) represented the most abundant phyla. In accordance with the relatively most abundant species found in our previous molecular study (9), *Pseudomonas oleovorans*, *Acinetobacter parvus*, and *Moraxella osloensis* were also detected here by cultivation in the door seal, while *Rhizobium radiobacter* was detected in the detergent drawer (Table 1) (9).

Many of the identified species represent environmental bacteria, typically found in water habitats or the human body, such as skin-associated bacteria. In addition, some of the identified species are well-known biofilm formers, such as *Staphylococcus epidermidis*, *Micrococcus luteus*, *Bacillus cereus* and *Pseudomonas sp.* (2, 4, 7, 9, 10, 19–22).

To estimate their pathogenic potential, the identified bacterial species were classified into biosafety risk groups (RG) (23). More than 50% (21 of 40 species) were affiliated with risk group 2 (RG 2) organisms, i.e. represent a potential health risk especially for immunocompromised patients, pregnant women, or elderly persons (12). 15 out of 21 identified RG 2 bacteria were found in the detergent drawer compartment (DD and DC), and 13 out of 21 RG 2 bacteria in the entire rubber door seal.

By far, micrococci and staphylococci were the most frequently isolated genera, which is in contrast to the different molecular studies mentioned here (9, 10) and might represent a cultivation bias. Micrococci and staphylococci represent ubiquitous microorganisms that are often isolated from skin and mucous membranes of humans and animals, but also from air and water. They grow fast under a broad range of cultivation conditions (24–26). However, they also have the ability of dormancy and might therefore well resist the dramatically changing environmental conditions inside washing machines (27, 28).

Staphylococci such as *S. epidermidis*, *S. lugdunensis*, *S. saprophyticus*, and *S. haemolyticus* possess a pathogenic potential and may also play a role in horizontal gene transfer of antibiotic resistance genes (29–31). The presence and transmission of such resistance genes throughout washing machines have already been confirmed for ß-lactamase (32). β-lactamase-producing *Klebsiella oxytoca* and *Klebsiella pneumoniae* species have also been isolated from washing machines before (11, 33). It can be speculated that these bacteria can be transferred to other surfaces, e.g. via bioaerosols (29–31). Notably, *Klebsiella oxytoca* was also found in our study, however without knowing its resistance pattern.

Besides bacteria, a few eukaryotic species were also isolated with the used cultivation conditions, all affiliated with *Ascomycota* (1 %). The most abundant genus was *Aspergillus*. *Aspergillus sp*. are saprophytic fungi and can recycle organic debris. *A. fumigatus* is a prevalent airborne fungal pathogen that can cause severe infections in immunocompromised people (34).

### Conclusions

Despite its small sample size, our study clearly shows that both detergent drawer and bottom door seal of domestic washing machines are significantly contaminated with cultivable bacteria, including significant shares of potentially pathogenic ones. Maximum loads can exceed 10^5^ CFU per cm^2^. For the sake of machine and laundry hygiene, both parts should be cleaned regularly. Markedly lower CFU counts from the top part of the door seal underline the importance of water for microbial contamination of washing machines. When not in use, machines should be let open to dry out. Bacterial species identified here and in molecular studies as quantitatively important for the washing machine microbiota represent test organisms with high practical relevance for antimicrobial efficacy testing.

## Funding

S.J. was funded by the German Federal Ministry of Education and Research (project WMP, grant number 13FH197PX6). The article processing charge was funded by the Baden-Württemberg Ministry of Science, Research and Culture and Furtwangen University in the funding programme Open Access Publishing.

## Acknowledgements

The authors wish to thank all volunteers who participated in the study and provided their washing machines for microbiological analyses.

## Conflicts of Interest

M.W. is affiliated with Henkel AG & Co. KGaA, a manufacturer of laundry and home care products. Henkel did not have any additional role in study design, data collection and analysis, decision to publish or preparation of the manuscript.

## References

1. Babič MN, Zalar P, Ženko B, Schroers H-J, Džeroski S, Gunde-Cimerman N. Candida and *Fusarium* species known as opportunistic human pathogens from customer-accessible parts of residential washing machines. Fungal Biol 2015; 119(2-3):95–113.

2. Novak Babič M, Gostinčar C, Gunde-Cimerman N. Microorganisms populating the water-related indoor biome. Appl Microbiol Biotechnol 2020; 104(15):6443–62.

3. Egert M. The BE-Microbiome-Communities with Relevance for Laundry and Home Care. SOFW J.:44–8.

4. Munk S, Johansen C, Stahnke LH, Adler-Nissen J. Microbial survival and odor in laundry. J Surfact Deterg 2001; 4(4):385–94.

5. Bloomfield SF, Exner M, Goroncy-Bermes P, Hartemann P, Heeg P, Ilschner C et al. Lesser-known or hidden reservoirs of infection and implications for adequate prevention strategies: Where to look and what to look for. GMS Hyg Infect Control 2015; 10:Doc04.

6. Gibson L. L., Rose J. B., Haas C. N. Use of quantitative microbial risk assessment for evaluation of the benefits of laundry sanitation. Am J Infect Control. 1999; 27(6):S34–S39.

7. Callewaert C, van Nevel S, Kerckhof FM, Granitsiotis MS, Boon N. Bacterial Exchange in Household Washing Machines. Front Microbiol. 2015; 6:1381.

8. Gattlen J, Amberg C, Zinn M, Mauclaire L. Biofilms isolated from washing machines from three continents and their tolerance to a standard detergent. Biofouling 2010; 26(8):873–82.

9. Jacksch S, Kaiser D, Weis S, Weide M, Ratering S, Schnell S et al. Influence of Sampling Site and other Environmental Factors on the Bacterial Community Composition of Domestic Washing Machines. Microorganisms 2019; 8(1).

10. Nix ID, Frontzek A., Bockmühl DP. Characterization of Microbial Communities in Household Washing Machines. TSD 2015; 52(6):432–40.

11. Schmithausen RM, Sib E, Exner M, Hack S, Rösing C, Ciorba P et al. The Washing Machine as a Reservoir for Transmission of Extended-Spectrum-Beta-Lactamase (CTX-M-15)-Producing *Klebsiella oxytoca* ST201 to Newborns. Appl Environ Microbiol 2019; 85(22).

12. Fritz B, Jenner A, Wahl S, Lappe C, Zehender A, Horn C et al. A view to a kill? - Ambient bacterial load of frames and lenses of spectacles and evaluation of different cleaning methods. PLoS ONE 2018; 13(11):e0207238.

13. R: A language and environment for statistical computing. Vienna, Austria: R Foundation for Statistical Computing; 2018. Available from: URL: https://www.R-project.org/.

14. RStudio: Integrated Development for R. Boston, MA: RStudio, Inc.; 2016. Available from: URL: http://www.rstudio.com/.

15. Wickham H, Sievert C. ggplot2: Elegant graphics for data analysis. Second edition. New York, NY, USA: Springer-Verlag; 2016. (Use R!).

16. Wickham H. Reshaping Data with the reshape Package. J. Stat. Soft. 2007; 21(12).

17. scales: Scale Functions for Visualization; 2018. Available from: URL: https://www.rdocumentation.org/packages/scales.

18. Stapleton K, Hill K, Day K, Perry JD, Dean JR. The potential impact of washing machines on laundry malodour generation. Lett Appl Microbiol 2013; 56(4):299–306.

19. Otto M. Staphylococcal biofilms. Curr Top Microbiol Immunol 2008; 322:207–28.

20. Vlamakis H, Chai Y, Beauregard P, Losick R, Kolter R. Sticking together: Building a biofilm the *Bacillus subtilis* way. Nat Rev Microbiol 2013; 11(3):157–68.

21. Matsuura K, Asano Y, Yamada A, Naruse K. Detection of *Micrococcus luteus* biofilm formation in microfluidic environments by pH measurement using an ion-sensitive field-effect transistor. Sensors (Basel) 2013; 13(2):2484–93.

22. Mann E. E., Wozniak D. J. Pseudomonas biofilm matrix composition and niche biology. FEMS Microbiol Rev. 2012; 36(4):893–916.

23. baua - Bundesanstalt für Arbeitsschutz und Arbeitsmedizin. TRBA 466: Einstufung von Prokaryonten (Bacteria und Archaea) in Risikogruppen; 2015 [cited 2020 Sep 9]. Available from: URL: https://www.baua.de/DE/Angebote/Rechtstexte-und-Technische-Regeln/Regelwerk/TRBA/TRBA-466.html.

24. Chiller K, Selkin BA, Murakawa GJ. Skin microflora and bacterial infections of the skin. J Investig Dermatol Symp Proc. 2001; 6(3):170–4.

25. Götz F, Bannerman T, Schleifer K-H. The Genera *Staphylococcus* and *Macrococcus*. In: Dworkin M, Falkow S, Rosenberg E, Schleifer K-H, Stackebrandt E, editors. The Prokaryotes: Volume 4: Bacteria: Firmicutes, Cyanobacteria. New York, NY: Springer US; 2006. p. 5–75.

26. Fang Z, Ouyang Z, Zheng H, Wang X, Hu L. Culturable airborne bacteria in outdoor environments in Beijing,China. Microb Ecol 2007; 54(3):487–96.

27. Kaprelyants AS, Kell DB. Dormancy in Stationary-Phase Cultures of *Micrococcus luteus*: Flow Cytometric Analysis of Starvation and Resuscitation. Appl Environ Microbiol 1993; 59(10):3187–96.

28. Cerca F, França Â, Pérez-Cabezas B, Carvalhais V, Ribeiro A, Azeredo J et al. Dormant bacteria within *Staphylococcus epidermidis* biofilms have low inflammatory properties and maintain tolerance to vancomycin and penicillin after entering planktonic growth. J Med Microbiol. 2014; 63(Pt 10):1274–83.

29. Madsen AM, Moslehi-Jenabian S, Islam MZ, Frankel M, Spilak M, Frederiksen MW. Concentrations of *Staphylococcus* species in indoor air as associated with other bacteria, season, relative humidity, air change rate, and *S. aureus*-positive occupants. Environ Res 2018; 160:282–91.

30. Kooken JM, Fox KF, Fox A. Characterization of *Micrococcus* strains isolated from indoor air. Mol Cell Probes 2012; 26(1):1–5.

31. Brandl H, Fricker-Feer C, Ziegler D, Mandal J, Stephan R, Lehner A. Distribution and identification of culturable airborne microorganisms in a Swiss milk processing facility. J Dairy Sci 2014; 97(1):240–6.

32. Rehberg L, Frontzek A, Melhus Å, Bockmühl DP. Prevalence of β-lactamase genes in domestic washing machines and dishwashers and the impact of laundering processes on antibiotic-resistant bacteria. J Appl Microbiol 2017; 123(6):1396–406.

33. Boonstra MB, Spijkerman DCM, Voor In ’t Holt, A. F., van der Laan, R. J., Bode LGM, van Vianen W et al. An outbreak of ST307 extended-spectrum beta-lactamase (ESBL)-producing *Klebsiella pneumoniae* in a rehabilitation center: An unusual source and route of transmission. Infect Control Hosp Epidemiol 2020; 41(1):31–6.

34. Latgé JP. *Aspergillus fumigatus* and aspergillosis. Clin Microbiol Rev 1999; 12(2):310–50.

